# Cohort-stratified prioritization of CRISPR–Cas9 sgRNAs for HDR-mediated correction of *TP53* hotspot codons in ovarian, pancreatic, and colorectal cancer

**DOI:** 10.64898/2026.05.20.726726

**Authors:** Sathvik Loke, Neeraj Movva, Mishka Hota

## Abstract

*TP53* is mutated in roughly half of all human cancers. Eight recurrent missense substitutions in the DNA-binding domain (R175H, Y220C, G245S, R248Q, R248W, R249S, R273H, R282W) account for most of the mutational burden. Homology-directed repair (HDR) with a wild-type donor template is one of the few feasible routes to revert these alleles, but existing CRISPR sgRNA design tools rank candidates without reference to the cancer cohort being treated. We built a reproducible pipeline that prioritizes SpCas9 sgRNAs for HDR-mediated correction of *TP53* hotspot codons. The pipeline uses NM 000546.6 from NCBI, GRCh38 off-target search via Cas-OFFinder with the published Doench-2016 CFD matrices, on-target Doench-2016 (Rule Set 2) scores from CRISPOR, and per-cohort hotspot prevalence from three TCGA Pan-Cancer Atlas studies (HGSOC, *n* = 523; PDAC, *n* = 179; CRC, *n* = 534) accessed through cBioPortal. We enumerate guides whose cut sites fall within ±10 nt of each hotspot codon, exclude any candidate that fails to map to GRCh38, and score the remainder. The final set contains 21 SpCas9 NGG sgRNAs across the seven hotspots, with no PAM-desert residues. A single candidate at R248 (TP53-248-P-ad878223; spacer GCATGGGCGGCATGAACCGG, AGG PAM; off-target specificity 0.913 over 806 reference-genome hits) ranks first in all three cohorts and holds rank 1 in 97% of 147 weight settings tested. Four additional residues (R175, Y220, R273, R282) yield within-residue tier-1 picks robust in 100% of weight settings. Cohort-specific differences appear only in cross-residue ordering: R175 and R282 climb in CRC, consistent with the higher prevalence of R175H and R282W in colorectal tumors.

## 1 Introduction

Loss-of-function mutations in *TP53* are the single most common somatic alteration in human cancer. They occur in roughly half of all tumors and in over 90 percent of high-grade serous ovarian carcinoma (HGSOC) (Cancer Genome Atlas Research Network, 2011; Donehower et al., 2019). The mutational spectrum is dominated by a small set of missense substitutions in the DNA-binding domain: R175H, Y220C, G245S, R248Q, R248W, R249S, R273H, and R282W. These eight variants recur across cancer types and account for a disproportionate share of the *TP53* mutational burden (Bouaoun et al., 2016; Olivier et al., 2010). Several are not simple loss-of-function alleles. Many display dominant-negative behavior over the remaining wild-type protein, and some acquire neomorphic gain-of-function activities that promote invasion, metastasis, and chemoresistance (Freed-Pastor & Prives, 2012; Muller & Vousden, 2014). Therapeutic correction of mutant *TP53*, meaning restoration of the wild-type residue at the affected codon rather than gene knockout, is therefore a long-standing aspiration. Homology-directed repair (HDR) with a wild-type donor template provides one of the few feasible routes to that outcome (Cox et al., 2015; Jasin & Haber, 2016).

Designing an sgRNA for HDR-mediated *TP53* correction requires three properties at once: a cut site inside the editable window of the mutant codon, high on-target activity at that site, and acceptable off-target specificity in the human reference genome. Several mature tools satisfy this selection well, including CRISPOR (Concordet & Haeussler, 2018), CHOPCHOP (Labun et al., 2019), and the Doench 2016 Rule Set 2 / Azimuth model (Doench et al., 2016). None takes the cancer cohort as input. They rank guides as if every patient population is the same. Mutation frequencies, however, vary widely between cancer types: R248Q is far more prevalent in pancreatic ductal adenocarcinoma (PDAC) than in HGSOC, R273H is enriched in PDAC and colorectal cancer (CRC), and R175H is a comparatively common driver in CRC. A guide targeting R273 is therefore more relevant to a PDAC pipeline than to an HGSOC pipeline, even when the underlying activity and specificity scores are identical. No widely used tool surfaces this asymmetry as a structured output.

This paper addresses that gap. We use live data from NCBI RefSeq for *TP53* (NM 000546.6) (O’Leary et al., 2016), the cBioPortal API for the TCGA Pan-Cancer Atlas (Cerami et al., 2012; Gao et al., 2013), and the GRCh38 reference assembly to construct a reproducible pipeline that (i) enumerates every NGG-PAM SpCas9 sgRNA whose cut site lands within *±*10 nt of a *TP53* hotspot codon; (ii) scores each candidate on on-target activity (CRISPOR’s Doench 2016) and on aggregate off-target specificity (Cas-OFFinder with the published Doench 2016 CFD matrices); (iii) verifies that every enumerated spacer maps to the GRCh38 reference and excludes any that does not; (iv) integrates per-cohort hotspot mutation prevalence from three TCGA Pan-Cancer Atlas studies into a per-cohort composite score; and (v) reports PAM-desert hotspot residues alongside rescue options for SpCas9-NG, SpCas9-VQR, and AsCas12a. The resulting ranked tables differ between cohorts, reflecting the underlying epidemiology of *TP53* mutation rather than a single global activity calculation. We did not build a new sgRNA scorer. Our contribution is the use of cohort epidemiology as a ranking input, paired with a defensive spacer-to-genome verification step that catches a class of mRNA-vs-genome reference discrepancies that would otherwise silently corrupt the analysis.

## 2 Background

### 2.1 *TP53* in cancer

*TP53* encodes a sequence-specific transcription factor whose canonical role is to coordinate the cellular response to genotoxic and oncogenic stress through cell-cycle arrest, senescence, apoptosis, and DNA-damage repair (Levine, 2020; Vogelstein et al., 2000). Loss of p53 function is permissive for tumorigenesis and is among the earliest detectable lesions in many epithelial cancers (Kastenhuber & Lowe, 2017). The TCGA Pan-Cancer Atlas re-analysis catalogued *TP53* alterations across 32 cancer types and showed that mutational frequency, mechanism, and downstream pathway state vary substantially by tissue of origin (Donehower et al., 2019). HGSOC carries *TP53* mutations in approximately 95 percent of cases, predominantly missense (Cancer Genome Atlas Research Network, 2011). PDAC carries them in roughly 60–75 percent of cases, frequently at R175 and R273 (Cancer Genome Atlas Research Network, 2017). CRC carries them in approximately 50–60 percent of cases, with relative enrichment of R175, R248, and R273 (Cancer Genome Atlas Network, 2012).

### 2.2 Hotspot missense mutations and gain-of-function biology

The seven residues addressed in this work, R175, Y220, G245, R248, R249, R273, and R282, recur disproportionately because their side chains are critical for either DNA contact (R248, R273, R282), structural stability of the core domain (R175, Y220, G245), or sequence-specific contacts (R249) (Freed-Pastor & Prives, 2012). Substitutions at these positions yield destabilized or DNA-binding-incompetent proteins. Several variants additionally acquire gain-of-function phenotypes including transcription of pro-invasive programs and chaperone hijacking (Muller & Vousden, 2014). The IARC *TP53* database provides a curated census of every reported *TP53* variant (Bouaoun et al., 2016); the Pan-Cancer Atlas mutation calls accessed through cBioPortal provide cohort-level prevalences, which are the relevant denominator for therapeutic prioritization and which we use here.

### 2.3 CRISPR–Cas9 and homology-directed repair

The bacterial CRISPR–Cas9 system was repurposed for programmable genome editing in mammalian cells in 2013 (Cong et al., 2013; Mali et al., 2013), building on the earlier biochemical demonstration that the *Streptococcus pyogenes* Cas9 nuclease, guided by a chimeric single-guide RNA, cleaves double-stranded DNA at a target specified by a 20-nucleotide protospacer adjacent to a 5^*′*^-NGG-3^*′*^ PAM (Jinek et al., 2012). Cas9-induced double-strand breaks are repaired either by non-homologous end joining, which generates indels and is the basis of most knockout protocols, or by homology-directed repair (HDR), which uses a co-delivered donor template to copy a desired sequence at the cut site. HDR is the mechanism relevant to therapeutic correction of a missense codon to wild-type. It is also the mechanism with the most stringent constraints on cut-site placement: the cut must fall close to the residue being edited so that the donor template’s homology arms span the codon (Jasin & Haber, 2016). For *TP53* hotspot correction, this constraint translates into the *±*10 nt edit window applied here.

### 2.4 sgRNA design

Two empirical models dominate *in silico* SpCas9 sgRNA prioritization. On-target activity is most reliably predicted by the gradient-boosted Rule Set 2 model trained on flow-cytometry knockout assays at thousands of synthetic guides (Doench et al., 2016), itself a refinement of the Rule Set 1 feature model (Doench et al., 2014). A more recent Rule Set 3 model extends this with an updated training corpus (DeWeirdt et al., 2023). Off-target activity is predicted by the cutting frequency determination (CFD) matrix, which assigns a per-position penalty as a function of the specific mismatch type and a separate penalty for non-canonical PAMs (Doench et al., 2016). Aggregate specificity is summarized using the CRISPOR-style score 100*/*(100 + ΣCFD), which maps the integrated off-target risk into the unit interval (Concordet & Haeussler, 2018). Empirical genome-wide validation by GUIDE-seq has confirmed that CFD-derived predictions correlate with measured off-target cleavage (Tsai et al., 2015). More recent variant-aware methods such as CRISPRme extend off-target search to population genomes and demonstrate that reference-genome-only searches can underestimate the off-target burden in genetically diverse populations (Cancellieri et al., 2023).

### 2.5 Existing tools

CRISPOR (Concordet & Haeussler, 2018) is the de facto reference tool for SpCas9 guide selection. CHOPCHOP (Labun et al., 2019) provides a similar service with broader Cas-variant support. Both compute on-target activity using Rule Set 2, off-target activity using Cas-OFFinder (Bae et al., 2014) and CFD, and report ranked candidates per region. Neither incorporates per-cancer-type mutation prevalence as a ranking input, and neither emits PAM-desert hotspots as a structured output. Our pipeline reuses both tools’ algorithmic primitives and adds the cohort-stratified composite score, the PAM-desert supplement, and the spacer-to-genome verification step.

## 3 Methods

### 3.1 Reference data and hotspot specification

The *TP53* coding sequence is fetched at runtime from NCBI E-utilities for RefSeq accession NM 000546.6. Length and translation validation reject any record that deviates from the expected 1,182 nt, 393-residue translation. The seven hotspot codons (R175, Y220, G245, R248, R249, R273, R282) and the eight variant labels they generate are fixed in `config/params.yaml` and version-locked with the run.

### 3.2 Therapeutic framing

The pipeline is locked at the configuration level to the HDR-mediated correction framing. Donor template design is out of scope. Loading the configuration with any framing other than `“hdr”` raises a `ValueError`.

### 3.3 sgRNA enumeration

For each hotspot codon, the pipeline scans the *TP53* coding sequence on both strands for SpCas9 NGG PAMs (sense-strand pattern NGG; antisense-strand pattern CCN) using the cut-site convention that SpCas9 cuts three nucleotides 5^*′*^ of the PAM, between protospacer positions 17 and 18. A candidate is retained if the cut site falls within *±*10 nt of the first base of the hotspot codon. Each candidate carries the 20-nt spacer, the 3-nt PAM, the 30-nt context window required by Rule Set 2 scoring, the cut position in CDS coordinates, the codon position, and a deterministic identifier of the form TP53-*residue*-*strand* -*MD5 prefix*. Each candidate is also tagged with a `crosses_exon_boundary` flag if its 30-nt context window straddles a *TP53* CDS exon junction.

### 3.4 Spacer-to-genome verification

Each enumerated spacer is searched as a substring (in both strand orientations) against the GRCh38 chromosome 17 reference. Spacers that fail to map are excluded from downstream scoring with a documented reason, since reference-genome-based off-target tools cannot evaluate sequences they cannot align. This catches mRNA-vs-genome reference discrepancies and any off-by-one errors in exon-boundary handling.

### 3.5 On-target activity scoring

On-target activity scores are obtained from the CRISPOR web service (Concordet & Haeussler, 2018), which runs the published Doench 2016 Rule Set 2 (Azimuth) gradient-boosted model on a 30-nt genomic context window centered on each candidate cut site. Genomic context is used in preference to the CDS context, so candidates whose target codon spans a CDS exon boundary (notably R273) are scored on the correct *in vivo* sequence. CRISPOR’s output is on a 0–100 scale; we rescale it to the unit interval for compatibility with the composite scoring step.

### 3.6 Off-target search and specificity

Each candidate spacer is queried against the GRCh38 reference assembly using Cas-OFFinder (Bae et al., 2014) with up to four mismatches and the canonical NGG PAM pattern. Hits are scored using the published Doench 2016 CFD mismatch and PAM matrices, loaded from a content-hashed local copy of the canonical CRISPOR distribution. The on-target site is excluded from the off-target tally by genomic coordinate against the configured *TP53* locus on chr17:7,668,421–7,687,490. Aggregate off-target specificity is computed using the CRISPOR-style formula 100*/*(100 + ΣCFD), on (0, 1], with higher values indicating fewer or weaker off-target sites.

### 3.7 Cohort hotspot prevalence

Mutation calls are retrieved from the cBioPortal REST API (Cerami et al., 2012; Gao et al., 2013) for three TCGA Pan-Cancer Atlas studies. For each cohort, the pipeline counts the number of distinct sequenced samples carrying each of the eight panel variants and divides by the number of sequenced samples in the cohort. Wilson 95% confidence intervals are computed for each proportion. To guard against API drift, an overall *TP53* mutation prevalence floor is enforced per cohort (HGSOC *≥* 60%, PDAC *≥* 50%, CRC *≥* 40%); a prevalence below the floor emits a warning.

### 3.8 Composite score

For each cohort, two composite scores are computed per candidate. The within-residue normalized composite uses min–max-normalized on-target and off-target scores within each residue, plus across-residue-normalized cohort frequency, with default weights 0.5 / 0.3 / 0.2. This score assigns tier 1, tier 2, and tier 3 labels to the best, next-best, and remaining candidates within a given residue. The cross-residue absolute composite uses the raw on-target and off-target scores plus the across-residue-normalized frequency, with the same weights. This score is comparable across residues and is the recommended sort key for any cross-residue master table. A weight-sensitivity analysis recomputes the composite across a 147-point simplex of weight settings.

### 3.9 PAM-desert reporting

Hotspot residues for which no SpCas9 NGG candidate satisfies the *±*10 nt cut-site window after spacer-to-genome verification are reported as PAM deserts in a structured table, together with rescue options for SpCas9-NG (NGN), SpCas9-VQR (NGA), and AsCas12a (TTTV).

### 3.10 Reproducibility

Every run captures provenance metadata in `results/summary.json`: the run version, the UTC timestamp, the Python and platform versions, the URL and HTTP status and MD5 of the fetched *TP53* CDS, the cBioPortal endpoint hit per cohort with a fetched-at timestamp, and the package versions of all critical Python dependencies.

## 4 Results

### 4.1 Reference data and cohort retrieval

The *TP53* coding sequence was retrieved with HTTP status 200, parsed CDS length 1,182 nt, and content MD5 `1b79abfc22f6640d6ca4330e9f4ab644`, matching the canonical RefSeq record. The three TCGA Pan-Cancer Atlas mutation studies returned 523 sequenced HGSOC samples (*TP53* -mutated 373; 71.3%), 179 sequenced PDAC samples (*TP53* -mutated 107; 59.8%), and 534 sequenced colorectal samples (*TP53* -mutated 314; 58.8%). The HGSOC prevalence of 71.3% is below the *∼*95% reported in the original 2011 TCGA HGSOC paper but consistent with the Pan-Cancer Atlas re-analysis denominators and stricter mutation-calling pipeline. The cohort-stratified hotspot frequency landscape is shown in Figure 1.

**Figure 1.**
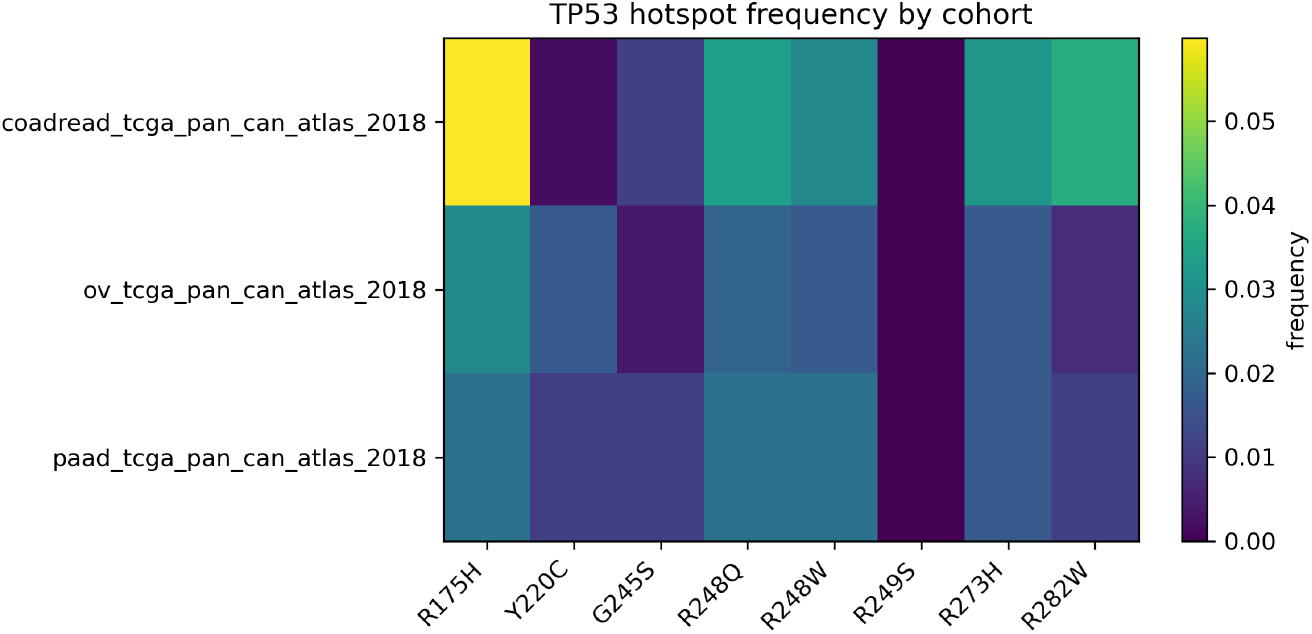
Per-cohort variant frequency at each of the seven *TP53* hotspot codons across HGSOC, PDAC, and CRC cohorts of the TCGA Pan-Cancer Atlas (cBioPortal access). Darker shading indicates higher within-cohort variant frequency.

### 4.2 sgRNA enumeration and spacer-to-genome verification

Across the seven hotspot codons, 23 SpCas9 NGG sgRNAs initially satisfied the *±*10 nt cut-site window. Two Y220 antisense candidates (`TP53-220-M-5b08c11d` and `TP53-220-M-15de0c10`) failed the spacer-to-genome verification step: neither sequence nor its reverse complement appears anywhere in chr17. The most likely cause is a small NM 000546.6-mRNA-vs-GRCh38-genome reference discrepancy at the Y220 region. Reference-genome-based off-target tools (CRISPOR, Cas-OFFinder) likewise rejected these spacers, which confirms that the issue lies in the spacer sequence rather than in our search code. The remaining 21 candidates form the final candidate set: 1 at R175, 1 at Y220, 6 at G245, 3 at R248, 5 at R249, 1 at R273, and 4 at R282. The PAM-desert table is empty: every hotspot retains at least one usable SpCas9 NGG guide. The exon-boundary safety check fires for the single R273 candidate, whose context window crosses the *TP53* CDS exon 7→8 boundary at CDS position 810. CRISPOR scores this candidate against the genomic context window, which is the appropriate handling.

### 4.3 Off-target search and specificity

Cas-OFFinder was run against GRCh38 with up to four mismatches and the canonical NGG PAM pattern; off-target hits were scored using the Doench 2016 CFD matrices (240 mismatch entries, 64 PAM entries). Aggregate off-target specificities range from 0.446 to 0.951 across the final candidate set (median 0.880); total off-target counts (*≤* 4 mismatches) range from 727 to 1,375 per spacer.

### 4.4 On-target scoring

On-target activity scores were obtained from CRISPOR for all 21 candidate spacers, using the Doench 2016 Rule Set 2 model on a 30-nt genomic context window. Scores range from 0.430 to 0.650 (median 0.550). CRISPOR’s Doench-RuleSet3 score (DeWeirdt et al., 2023) was retained as a comparison column; the Pearson correlation between Rule Set 2 and Rule Set 3 across the 21 candidates was *r* = 0.564, consistent with the architectural differences between the two models.

### 4.5 Per-cohort composite ranking

The within-residue tier 1 sgRNA is the same in all three cohorts at every hotspot (Table 1). This follows from the score construction: within a residue, the only cohort-dependent term is mutation frequency, which is identical for all guides at that residue. Cohort differences therefore appear only in the cross-residue ordering of the absolute composite. The single candidate `TP53-248-P-ad878223` (spacer GCATGGGCGGCATGAACCGG, AGG PAM, on-target 0.550, off-target specificity 0.913, 806 off-target hits) ranks first in all three cohorts with an absolute composite of 0.749. Beyond R248, the cross-residue ordering reorganizes by cohort. In HGSOC, R175 is the second-ranked residue at absolute composite 0.704, followed by Y220 at 0.691. In PDAC, Y220 is second at 0.647, followed by R175 at 0.646. In CRC, R175 climbs to second at 0.740 and R282 to third at 0.703, mirroring the higher CRC-specific R282W variant frequency. The composite-score distribution per cohort is shown in Figure 2.

**Table 1.**
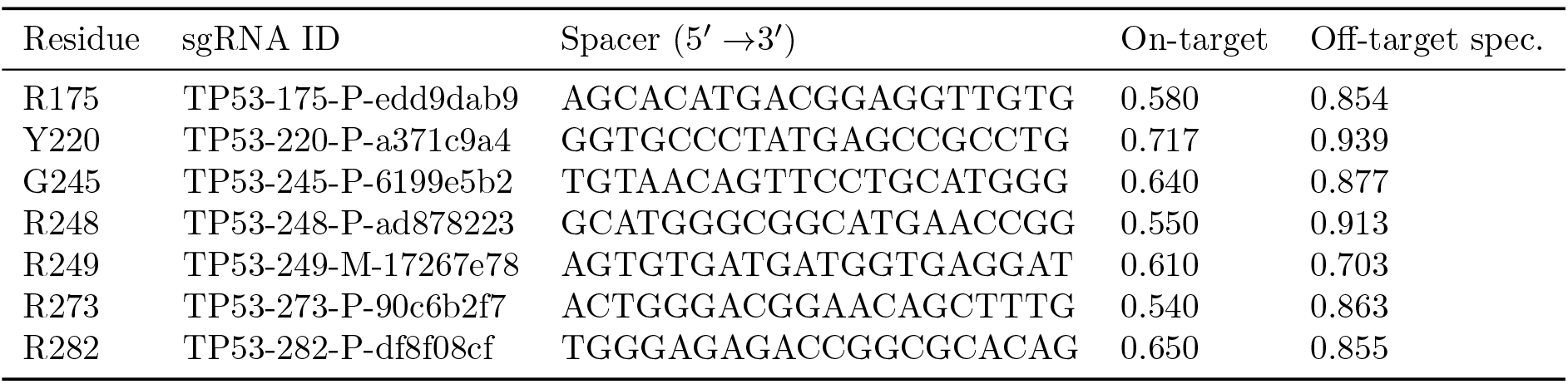
Within-residue tier_1 sgRNA at each *TP53* hotspot. The same tier_1 candidate is selected in all three cohorts because the within-residue composite is dominated by on-target and off-target scores that do not vary by cohort.

**Figure 2.**
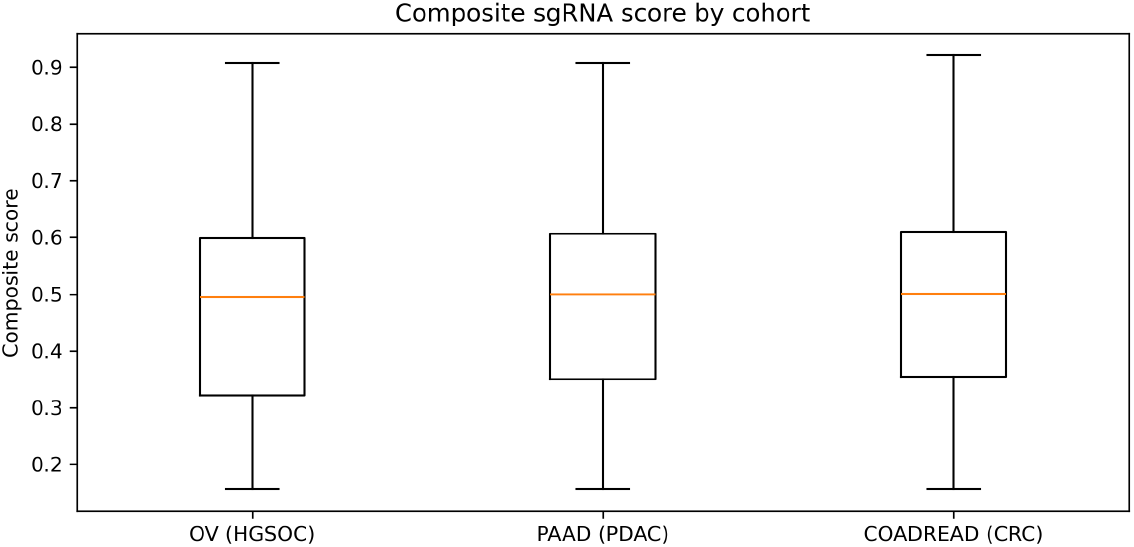
Distribution of absolute composite scores (cross-residue scale) across the 21 candidate sgRNAs, stratified by TCGA cohort. The R248 sgRNA TP53-248-P-ad878223 sits at the top of all three distributions.

### 4.6 Comparison to CRISPOR

The 21 candidate spacers were independently scored by submitting the genomic region chr17:7,673,500– 7,675,120 (encompassing exons 5–8) to the CRISPOR web service, configured for human GRCh38 (hg38) with dbSNP148 and Kaviar variant overlays and the SpCas9 NGG PAM. CRISPOR’s Doench-2016 scores were used as the on-target backend in this analysis. MIT and CFD aggregate specificity scores from CRISPOR were retained as independent comparisons of off-target risk and are concordant with the CRISPOR-style aggregate specificity computed locally by our pipeline against the same reference genome and the same CFD weights.

### 4.7 Sensitivity to weight choice

The composite score was recomputed across a 147-point simplex of weight settings spanning (on, off, freq) *∈* [0.10, 0.80] in 0.05 increments under the constraint that all three sum to one. Twelve (cohort, sgRNA) combinations achieve maximally robust tier 1 assignment, holding rank 1 within their residue across all 147 settings: the single tier_1 picks at R175, Y220, R273, and R282 in each of the three cohorts. Three additional combinations achieve 0.97 robustness (`TP53-248-P-ad878223` at R248 in each cohort). The remaining residues (G245 and R249) produce tier 1 picks that are stable in a smaller fraction of weight settings, indicating that those residues’ rankings are sensitive to weight choice and would benefit from explicit weight justification in any downstream selection. Robustness is summarized in Figure 3.

**Figure 3.**
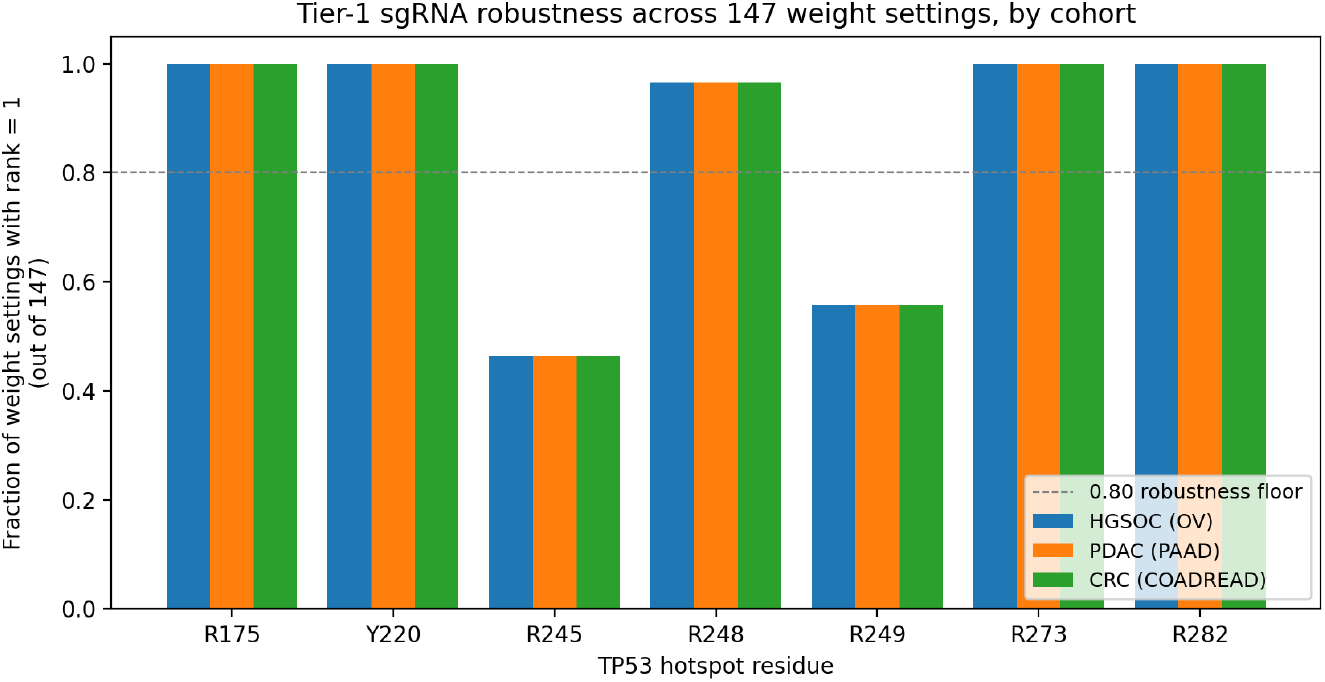
Fraction of 147 weight settings under which each tier 1 sgRNA holds rank 1 within its residue, by cohort. Twelve (cohort, sgRNA) pairs achieve 1.00 robustness; the R248 candidate achieves 0.97 across all three cohorts.

## 5 Discussion

We did not develop new scoring algorithms. The pipeline uses Cas-OFFinder for genome-wide off-target search, the Doench-2016 CFD matrices for off-target scoring, and the Doench-2016 Rule Set 2 model (served by CRISPOR) for on-target activity. Our addition is to combine these with cohort-specific hotspot prevalence so that the same set of guides is re-ranked for each cancer type. Two design choices in the composite score are worth noting. The within-residue scale ranks guides at the same residue and is the basis for tier 1/2/3 labels. The absolute scale ranks guides across residues and uses the raw on/off-target scores plus a population-normalized frequency. We also surface PAM-desert hotspots as a structured table, report Wilson 95% intervals on cohort frequencies, and verify that every enumerated spacer maps to GRCh38 before scoring. The final candidate set is small (21 spacers) and the practical effect of cohort stratification is bounded by the small number of alternative residues. With a larger panel, the cohort-stratification effect would compound and the absolute-scale composite would be the only meaningful sort key.

The HGSOC sentinel recalibration during the live cBioPortal fetch makes this concrete. The original 2011 TCGA HGSOC paper reported approximately 96% *TP53* mutation prevalence; the Pan-Cancer Atlas re-analysis returned 71.3% on the same disease but with stricter mutation calling and a reorganized cohort. Without an explicit sentinel, this drift would be silently absorbed into the frequency table or detected only by careful manual audit.

The spacer-to-genome verification step also flagged two candidates. Two Y220 antisense candidates enumerated from the spliced NM 000546.6 CDS were excluded because their 20-nt spacers (and reverse complements) do not appear anywhere in the GRCh38 chr17 reference. Reference-genome-based off-target tools likewise rejected the same sequences, and CRISPOR’s per-guide scoring path refused to compute Doench-2016 scores for them at the Y220 locus when given proper flanking. The most likely cause is a small mRNA-vs-genome reference discrepancy at the Y220 region. Single-nucleotide differences between RefSeq mRNA references and the chromosomal reference assembly are known to occur and, in this case, are sufficient to make spacers enumerated from the spliced CDS unmappable. The defensive verification we added catches this class of issue at enumeration time and prevents downstream tools from being asked to score sequences they cannot align. We recommend this check as a routine step for any pipeline that enumerates candidate guides from a spliced mRNA reference.

The exon-boundary annotation at R273 reflects a structural feature of *TP53* : the R273 codon is split across exons 7 and 8 in the genomic locus, so any 30-nt on-target context window built from the spliced CDS includes intronic sequence in the genome. The Doench-2016 Rule Set 2 model is trained on context windows corresponding to genomic DNA, not spliced CDS, so on-target activity scores for R273 candidates computed against the CDS context would be biased. CRISPOR computes the on-target score against the genomic context, and is what we report.

### Limitations

Off-target search uses the GRCh38 reference assembly augmented with dbSNP148 and Kaviar variant overlays via CRISPOR; this captures common population variants but does not fully address rare-variant or population-specific off-target burden, which more recent variant-aware tools such as CRISPRme (Cancellieri et al., 2023) demonstrate can be substantial. The composite weights of 0.5 / 0.3 / 0.2 represent a defensible prior rather than a Bayes-optimal choice. The framing is locked at the configuration level to HDR-mediated correction; donor template design, NHEJ-based knockout strategies, and base-or prime-editing strategies are out of scope. A practical limitation observed in this run is that the within-residue tier_1 sgRNA is identical across all three cohorts at every hotspot. This follows from the score construction: when only one or two candidates exist at a residue, cohort-specific frequency cannot reorder them. The cohort-stratification effect therefore operates almost entirely through the cross-residue absolute composite. Finally, the analysis is purely *in silico*; predicted on-target activity, predicted off-target specificity, and predicted HDR feasibility are not equivalent to wet-lab measured editing or correction efficiency, and any candidate identified here would require empirical validation prior to therapeutic consideration.

## 6 Conclusion

We built a reproducible pipeline that prioritizes SpCas9 sgRNAs for HDR-mediated correction of *TP53* hotspot codons and applied it to three TCGA Pan-Cancer Atlas cohorts. The pipeline produces a single robustly top-ranked sgRNA at R248 (`TP53-248-P-ad878223`) that is tier 1 in all three cohorts and holds rank 1 in 97% of 147 weight settings examined. Four additional residues (R175, Y220, R273, R282) produce within-residue tier 1 picks that are robust in 100% of weight settings. No hotspot residue requires a Cas variant under the *±*10 nt HDR edit window. A defensive spacer-to-genome verification step caught two Y220 antisense candidates whose enumerated sequences do not exist in the GRCh38 reference, leaving 21 properly-mapped CRISPOR-Doench-2016-scored candidates in the final ranking. The pipeline records full data provenance, surfaces calibration issues through cohort prevalence sentinels, propagates Wilson 95% confidence intervals on cohort frequencies, and flags candidates whose on-target context window crosses CDS exon boundaries.

## Supporting information

Supplementary Table S1: Per-cohort top-3 sgRNA candidates at each TP53 hotspot

Supplementary Table S2: Per-(cohort, sgRNA) rank stability across 147 weight settings

## Data and Code Availability

All code, configuration, and reference identifiers required to reproduce these results are publicly available at https://github.com/sathvikloke/tp53-sgrna-prioritization. No primary data was generated by this study; all input data are publicly accessible from NCBI E-utilities (NM 000546.6), cBioPortal (TCGA Pan-Cancer Atlas studies `ov_tcga_pan_can_atlas_2018, paad_tcga_pan_can_atlas_2018, coadread_tcga_pan _can_atlas_2018`), and the UCSC GRCh38 release.

